# Hierarchical learning of statistical regularities over multiple timescales of sound sequence processing: A dynamic causal modelling study

**DOI:** 10.1101/768846

**Authors:** Kaitlin Fitzgerald, Ryszard Auksztulewicz, Alexander Provost, Bryan Paton, Zachary Howard, Juanita Todd

## Abstract

The nervous system is endowed with predictive capabilities, updating neural activity to reflect recent stimulus statistics in a manner which optimises processing of expected future states. This process has previously been formulated within a predictive coding framework, where sensory input is either “explained away” by accurate top-down predictions, or leads to a salient prediction error which triggers an update to the existing prediction when inaccurate. However, exactly how the brain optimises predictive processes in the stochastic and multi-faceted real-world environment remains unclear. Auditory evoked potentials have proven a useful measure of monitoring unsupervised learning of patterning in sound sequences through modulations of the mismatch negativity component which is associated with “change detection” and widely used as a proxy for indexing learnt regularities. Here we used dynamic causal modelling to analyse scalp-recorded auditory evoked potentials collected during presentation of sound sequences consisting of multiple, nested regularities and extend on previous observations of pattern learning restricted to the scalp level or based on single-outcome events. Patterns included the regular characteristics of the two tones presented, consistency in their relative probabilities as either common standard (*p* = .875) or rare deviant (*p* = .125), and the regular rate at which these tone probabilities alternated. Significant changes in connectivity reflecting a drop in the precision of prediction errors based on learnt patterns were observed at three points in the sound sequence, corresponding to the three hierarchical levels of nested regularities: (1) when an unexpected “deviant” sound was encountered; (2) when the probabilities of the two tonal states altered; and (3) when there was a change in rate at which probabilities in tonal state changed. These observations provide further evidence of simultaneous pattern learning over multiple timescales, reflected through changes in neural activity below the scalp.

**Author summary:** Our physical environment is comprised of regularities which give structure to our world. This consistency provides the basis for experiential learning, where we can increasingly master our interactions with our surroundings based on prior experience. This type of learning also guides how we sense and perceive the world. The sensory system is known to reduce responses to regular and predictable patterns of input, and conserve neural resources for processing input which is new and unexpected. Temporal pattern learning is particularly important for auditory processing, in disentangling overlapping sound streams and deciphering the information value of sound. For example, understanding human language requires an exquisite sensitivity to the rhythm and tempo of speech sounds. Here we elucidate the sensitivity of the auditory system to concurrent temporal patterning during a sound sequence consisting of nested patterns over three timescales. We used dynamic causal modelling to demonstrate that the auditory system monitors short, intermediate and longer-timescale patterns in sound simultaneously. We also show that these timescales are each represented by distinct connections between different brain areas. These findings support complex interactions between different areas of the brain as responsible for the ability to learn sophisticated patterns in sound even without conscious attention.

## Introduction

The alignment of neural activity to reflect recent stimulus statistics is a fundamental feature of the nervous system. Exponential reductions in neural firing with stimulus repetition form the physiological basis for a range of neural processes including sensory adaptation (1–4), associative learning (5, 6), and simple change detection (7, 8). Empirical and theoretical studies also suggest that predictive properties extend beyond single neurons and are applied with greater complexity throughout neural networks to actively generate inferences about future states in a manner consistent with Bayesian learning (9–13). Learnt causal relationships between stimuli and the structure of the environment are adaptive; they permit the pre-selection of adaptive behaviour and conserve processing resources for predicted events (14). Yet, the mechanisms by which the brain can optimise these associations in a complex and ever-changing natural environment remain unclear.

The natural environment comprises a multitude of regularities which constantly change and do so at different rates, with varying degrees of reliability. The brain is assumed capable of differentiating these states through a temporal hierarchy where different brain regions are sensitive to representing dynamics at different temporal scales. At the lowest level, sensory cortices encode fast-timescale dynamics underlying simple sensory processing, whilst the highest level involves the prefrontal cortex engaging the more complex functions required to represent slower-changing environmental states such as consistent variability in a given context (15, 16). At the neural level, individual neurons have time constants on the scale of milliseconds, post-synaptic gain control modulates precision on the scale of tens or hundreds of milliseconds, whilst connection strengths encode causal regularities that emerge more slowly (17). Computational models incorporating these hierarchical dependencies have been shown to predict actual neural responses and behaviour with a good degree of accuracy, and provide a suitable framework for hierarchical learning over multiple temporal scales (18–20). These hierarchical generative models of predictive coding assume that each neural population must reconcile existing predictions about input from the higher/more frontal level above with sensory input from the lower/more temporal level below, resulting in current input either being “explained away”, or a “prediction error” which drives for an update to the existing prediction. At each point these prediction errors are weighted by precision, or the strength and reliability of predictions and input, which determines the rate of new learning or readiness to update predictions accordingly (21, 22). Precision weighting is assumed to be implemented as neural gain modulation mediated by classical neuromodulators and N-methyl d aspartate (NMDA) dependent plasticity (23), however empirical data confirming the neurophysiology which supports this complex learning is limited.

Auditory evoked potentials (AEPs) provide a mode to study predictive processes within an implicit learning framework. AEPs are automatic, non-invasive and easily translated to populations including infants, clinical groups and the elderly. The N2a or mismatch negativity (MMN) is a negative deflection in AEP amplitude which emerges when comparing the response to an unexpected or low-probability sound to that of an expected, repetitive or high-probability sound (24–26). MMN increases in magnitude with the degree of deviance and is therefore considered an indicator of relative “surprise” (27). Modulations of MMN amplitude have been used as a proxy for surprise in a breadth of studies of perceptual inference and learning, including as evidence for hierarchical learning processes (e.g., 28– 30). However, the majority of these studies have focused on single-trial MMNs elicited following a simple deviation from a local pattern only (e.g., the traditional oddball paradigm and roving paradigms), and there is limited research into the impact of deviance occurring in a broader statistical context. More recently, growing evidence of the impact of varying degrees of uncertainty on precision weighting has been empirically shown via systematic modulations of the MMN observed in scalp-recorded AEPs during a multiple-timescale paradigm (31–35).

The multiple-timescale paradigm has used AEPs to reveal hierarchical learning during sound sequences consisting of multiple nested temporal regularities to create varying degrees of surprise. Early studies of this kind employed two tones which were presented with either standard (*p* = .875) or deviant probability (*p* = .125; local surprise), and alternated in these roles at a regular rate of every 0.8 minutes in “unstable” sequences or every 2.4 mins in “stable” sequences to create an additional, intermediate level of surprise when the relative probabilities of the two tones suddenly change (31,32,36,37). Despite local equivalence in sound probability ratios between standard and deviant across the two block types overall, sequences designed in this way have demonstrated AEP data consistent with a primacy effect, seen as higher precision in the prediction models for blocks consistent with how the sequence begins (large MMN throughout for the original block) relative to the blocks that represent the alternate probabilities (small MMN initially that increases with local stability within reversed blocks; (35,36,38). This differential precision is likened to lower and higher levels of expected uncertainty respectively, derived from an estimate of the volatility (conditional variance) of the current environment (39). In the absence of any existing priors, the model for the original block type is thought to be formed with high learning rates and high precision as the probabilities are rapidly used to predict the sound environment. The level of uncertainty drops rapidly and significantly over time as the model proves effective in predicting this context. In contrast, the model associated with the reversed block type develops in response to a gross violation in the existing high-precision model (which by then is associated with low expected uncertainty) when the initial deviant begins to repeat triggering a series of prediction errors. In this respect, this second context may be associated with a higher level of estimated volatility in the environment, as the transition to this context represents a substantial contextual change and learning rates are elevated accordingly. This error frequency is associated with a high level of surprise triggering a drop in model precision and an elevation in uncertainty whilst the internal model is updated.

The most recent iteration of the multiple-timescale paradigm introduced a third level of patterning within the sound sequences to investigate the sensitivity of the perceptual-cognitive system to regularities which unfold over even longer timescales, and the impact on learning when these higher order patterns are violated (34; see also 35). In this study, the previously mentioned “stable” and “unstable” sequences were concatenated in order to introduce a third level of “superordinate” surprise when the regular rate of alternation in tone tendency changes during a sequence by transitioning from “stable” sequence components comprised of 2.4-minute blocks to relatively more unstable sequence components comprised of 0.8-minute blocks, or vice versa. This modification resulted in the presentation of an “increasing-stability” sequence followed by a “decreasing-stability” sequence as represented in Figure 4 (see Materials and Methods).

Specific patterns of AEP modulation observed in this study showed that block type (intermediate-level predictability) remained influential until block length regularity was broken. When the original block type violated block length predictions (i.e., either by changing sooner or later than expected), MMN amplitude to this “first deviant” decreased, likely explained by a significant drop in precision due to superordinate surprise (34; see also 35). In contrast, MMN amplitude to deviants in the alternate block type (i.e., “second deviants”) were unaffected by the block length violation.

Whilst multiple-timescale studies have been informative in revealing the sensitivity to hierarchical patterning in sound through distinct modulations of scalp-recorded MMN, it remains unclear exactly where in the temporal hierarchy of the brain these changes occur due to the poor spatial acuity of sensor-level analyses and their limited focus on select electrodes, latencies and components. Here we used dynamic causal modelling (DCM) as an alternative method which explains the entire time-course and scalp topography of these data in terms of neurobiologically plausible mechanisms at the source level (i.e., arising from interactions between neural populations within and between sources; 40,41). This method provides a plausible mechanistic explanation of the observed data which cannot be offered by AEP alone, and therefore has greater sensitivity to test assumptions about underlying neurophysiology. DCM is also shown to have a greater sensitivity than neural data to variance in the population which is useful in the investigation of clinical groups (42).

The present study will apply DCM to data from the most recent multiple-timescale study by Fitzgerald and Todd (34) with the expectation that we will see evidence of differential modulation of connectivity and precision associated with different levels of surprise (local, intermediate, and superordinate). Timescale effects will be modelled using a DCM of a six-source hierarchical network consisting of sources in bilateral primary auditory cortex (A1), superior temportal gyri (STG) and inferior frontal gyri (IFG) given their best model evidence in previous DCM studies of the MMN (43–46). We expect to see the impact of pattern violations expressed differently in the network in a hierarchical manner dependent on the relative timescale of violation. More specifically we hypothesize that predictions/prediction errors will lead to connectivity change at hierarchically lower levels for violations of short timescales, and at higher levels for violations of long timescales. In doing so we seek to further establish the ability of the sensory system to perform sensory learning within unstable and oft-changing environments, and the underlying neural network which is employed in service of this aim.

## Results

### Sensor space results

The sensor space results derived from the common-average referenced AEPs replicated those observed by Fitzgerald and Todd (34), and a detailed outline is provided in S1 Appendix. Pertinent to the current DCM analysis is the observation that MMN to the first deviant (60 ms) showed clear modulation across the sequence where it was consistently smaller after a change in block length for decreasing stability (*t*(18) = −2.52, *p* < .05, corrected) and increasing stability (*t*(18) = 2.72, *p* < .05, corrected), with no significant change in MMN amplitude to the second deviant (30 ms) for either conditions. Given that DCM captures the entire epoch, analysis of mean amplitude of the P3 component was also conducted and revealed that P3 was significantly larger (more positive) for the 60 ms tone after a superordinate structure violation for the decreasing stability sequence only *t*(18) = - 2.50, *p* < .05, corrected).

Two-tailed t-tests of standard and deviant AEPs (per (62); *p* < .05, corrected) also replicated the finding that order-driven effects on MMN amplitude were evident in the deviant AEP, and significant differences in the deviant AEP between the two sequences were observed for the 60 ms tone only (i.e, the tone that was heard first as a local deviant), confirming the apparent insensitivity of the 30 ms deviant (i.e., the tone that was heard first as a local standard) AEP to order effects. The confinement of this differential sensitivity in responses to the two tones to the first-deviant AEP specifically emphasises order-driven effects in difference waveforms as related to a difference specifically in how these two contexts are treated, an assertion we aimed to assess using DCM.

### Connectivity effects

The effects of local deviance (standard vs deviant), superordinate (heard-first vs heard second) deviance and their interaction were first modelled separately for the 30 ms and 60 ms tone, whilst the effects of intermediate deviance (second/30 ms vs first/60 ms deviant) superordinate deviance (heard-first vs heard second) and their interaction were modelled on both tones together for deviant responses only in the second analysis. In both analyses, effects on connectivity were modelled within a six-source cortical network comprised of bilateral sources A1, STG and IFG. This choice was motivated by the goal of elucidating the hypothesised rostro-caudal temporal hierarchy in the brain where lower and higher levels are differentially sensitive to prediction errors at shorter and longer timescales respectively (18, 28). This specific selection of nodes is also in accordance with the sources chosen in previous DCM analyses of auditory MMN paradigms (43, 44). The full model permitted changes in ascending, descending, and intrinsic coupling between sources (model FBi in Figure 1), and was compared with a set of reduced models consisting of changes in each parameter alone, each combination, and a null model permitting no changes, resulting in a total of 8 x 8 models for comparison (see Figure 1 for representation of full model space).

**Figure 1.**
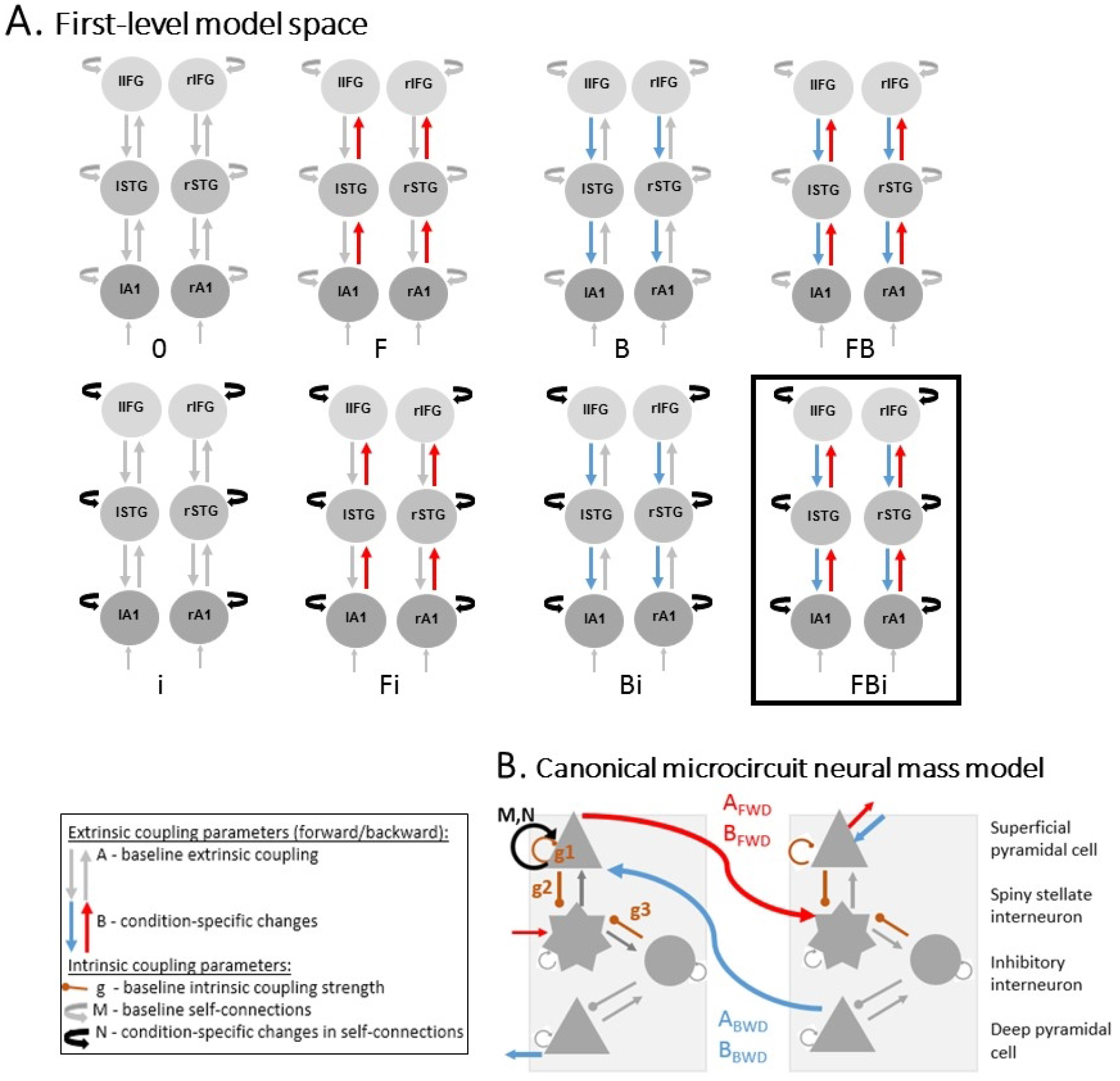
Representation of model space. Effects of local, intermediate and superordinate deviance were modelled within a six-source coupled network comprised of bilateral A1, STG and IFG, and permitting changes in ascending, descending and intrinsic connectivity. The same subset of parameters was allowed to vary for all factors resulting in a total of 8 discrete models shown in Figure 2A. Winning model FBi is indicated by a square outline. Each source within the model space was represented according to the canonical microcircuit neural mass model shown in Figure 2B, where intrinsic coupling parameters estimate the connectivity between superficial pyramidal cells, spiny stellate interneurons, inhibitory interneurons and deep pyramidal cells which contribute to condition-specific changes in extrinsic connectivity between sources.

BMR at the group-level was performed on all 64 models. The overall winning model was the most complex model permitting changes in ascending, descending and intrinsic connections (i.e., model FBi; see Figure 1A). This model was favoured in 100% of individual subjects in all analyses (30 ms and 60 ms tone modelled separately and modelled together), with a posterior probability exceeding 0.99 in all cases.

### Local deviance – Deviant relative to standard

Bayesian parameter averages for each connection type demonstrated similar directions of changes in connectivity for both the 30 ms and 60 ms tones when encountered as a local deviant relative to when encountered as a local standard. The direction and magnitude of these significant connectivity changes are displayed in Figure 2, and a full summary of parameter averages is provided in Table S1 and Table S2. Local deviance was associated with an increase in connection strength in all ascending connections for both tones, consistent with increased prediction error signalling. Local deviance was similarly associated with an asymmetrical change in intrinsic connectivity at A1 characterised by decreased intrinsic feedback at left A1 and increased intrinsic feedback at right A1 reflecting changes in the inhibitory self-suppression of prediction error at each of these sources when encountered as a deviant relative to standard. The significant (*p* < .005) changes in intrinsic connectivity were more widespread across the hierarchy for the second deviant (30 ms) than first deviant (60 ms) tone, with a decreased self-inhibition at bilateral STG and increased self-inhibition at rIFG for the 30 ms tone only reflecting increased and decreased gain of prediction error signalling at each of these sources, respectively. There was a specific unilateral increase in the strength of descending connections from STG to A1 which differed in location for the two tones, occurring from left STG to A1 for a 30 ms deviant, and right STG to A1 for a 60 ms deviant.

**Figure 2.**
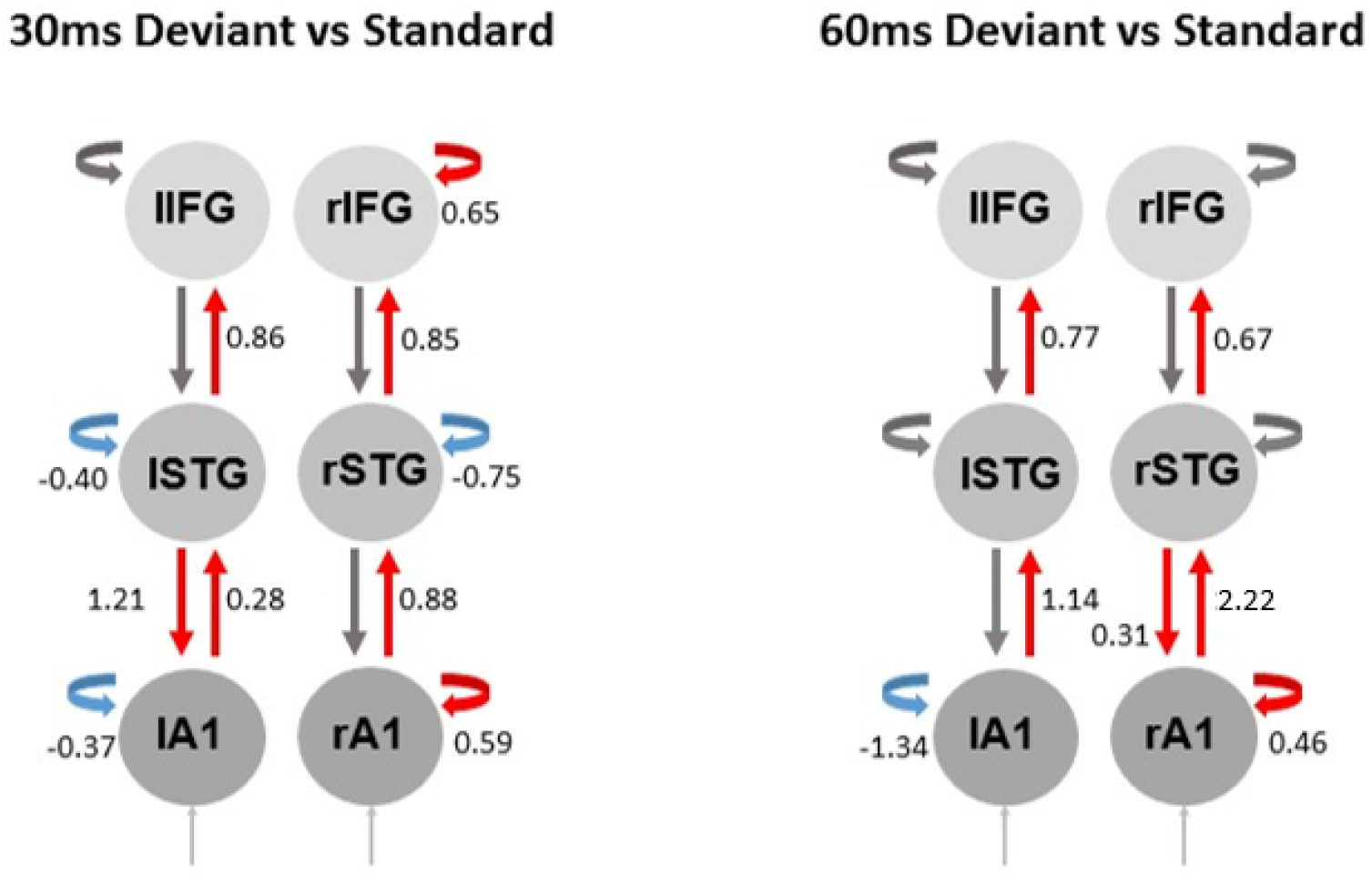
Connectivity changes associated with local deviance. Coloured arrows indicate significant increases (red) and decreases (blue) in ascending (upward arrows), descending (downward arrows) and intrinsic (curved arrows) connectivity strength associated with a 30 ms (left panel) and 60 ms (right panel) tone when encountered with deviant probability (*p* = .125) as relative to the same tone encountered with standard probability (*p* = .875). Values represent magnitude of difference in Bayesian parameter estimates relating to the significant modulation of extrinsic connections (B parameters; ascending/descending) and activity-dependent effects on intrinsic connectivity (N; within-source). Significance was assessed against a critical value of *p* < 0.005 using the procedure outlined in the Method.

### Higher-order effects – First vs Second deviant; Before versus after a block length change

Given that sensor space analyses revealed the differences in MMN to be driven by modulation of evoked responses to the tones as deviants as in previous multiple timescale MMN studies (e.g., 32,34–36,38), the impact of higher-order changes on the inferential network was assessed in a separate DCM analysis comprised of deviant AEPs only. Planned contrasts between deviant responses to the 30 ms and 60 ms tones, the two sequence components (before and after a change in superordinate structure) and their interaction were conducted to test for significant differences in connectivity associated with an order-driven modulation based on initial tone roles and with a superordinate pattern violation. This analysis revealed significant (*p* < .005) interactions throughout the network whereby changes related to superordinate sequence structure had differential impacts on network connectivity dependent on whether the deviant was a 60 ms or 30 ms tone. This differential effect is visually apparent in the plots of estimated parameter changes for the two tones over the entire sound sequence and specifically after a superordinate change, as displayed in Figure 3. A full summary of parameter averages is provided in Table S3 and Table S4.

**Figure 3.**
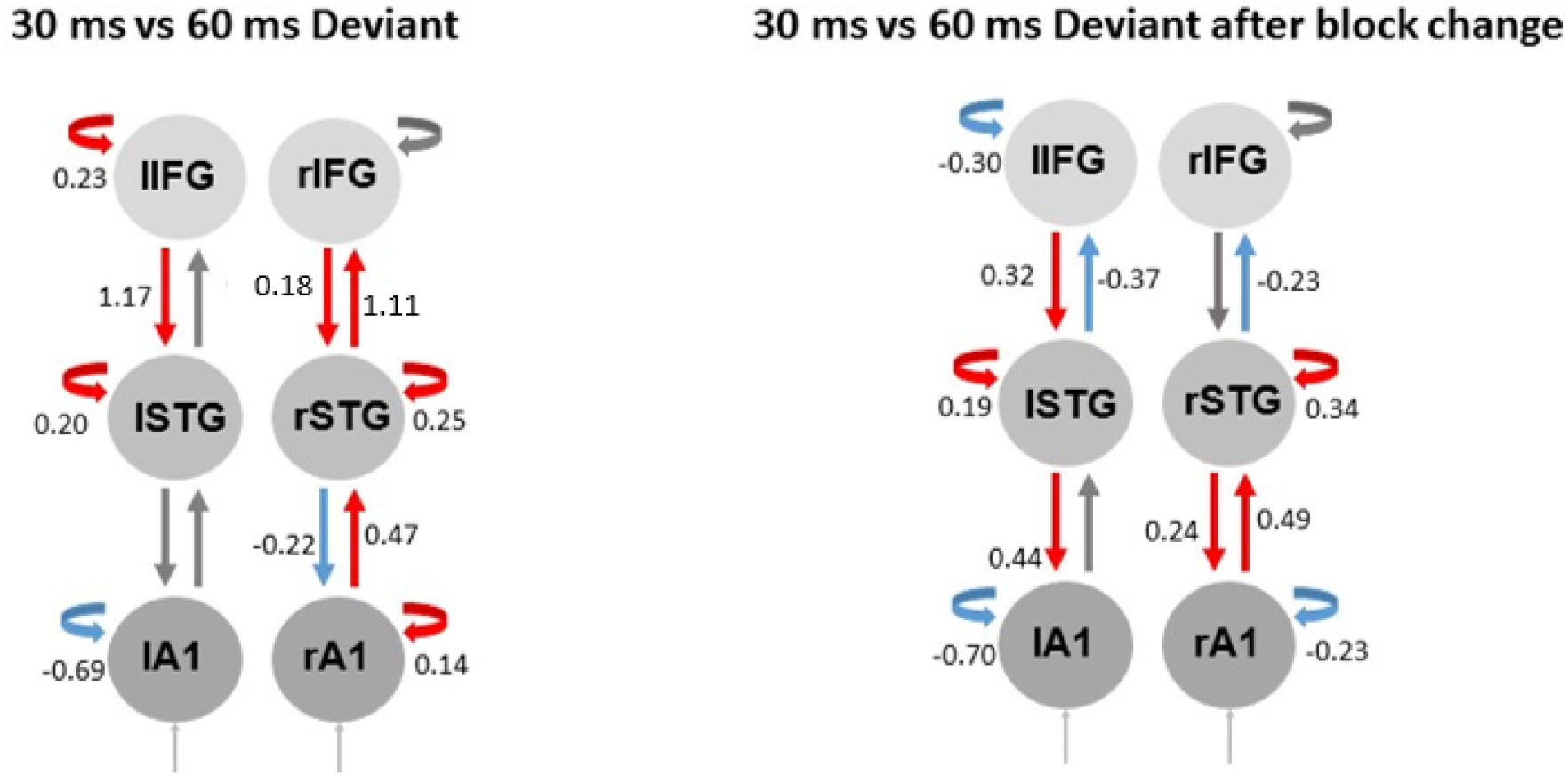
Connectivity changes associated with intermediate and superordinate deviance. Estimated changes in Bayesian connectivity parameters associated with a 30ms tone as deviant compared to a 60 ms deviant in general, reflecting intermediate deviance (left panel), and significant changes in connectivity for a 60 ms deviant as compared to a 30ms deviant specifically after a change in superordinate sequence structure (right panel). Coloured arrows indicate significant increases (red) and decreases (blue) in ascending (upward arrows), descending (downward arrows) and intrinsic (curved arrows) connectivity strength. Values represent magnitude of difference in Bayesian parameter estimates. A critical value of *p* < 0.005 was applied for significance testing using the procedure outlined in the Method.

First, significant differences in connectivity dynamics were observed when comparing the coupling changes associated with the deviant in the second block context (30 ms tone) compared to the deviant in the first block context (60 ms tone) across the overall sequence (Figure 3, left). The 30 ms tone, which was always the second deviant and therefore the initially repetitive standard, was associated with a significantly higher level of increased ascending error signalling, and a significantly higher level of increased inhibitory gain throughout most of the hierarchy (with the exception of lA1 and rIFG) compared to the 60 ms tone. The 30 ms tone was also associated with a higher level of increased descending message passing from higher levels (IFG), and comparatively lower descending message passing from lower levels (rSTG to rA1) relative to the 60 ms tone as deviant. In general, the lower precision of 30 ms prediction errors was therefore associated with higher forward coupling change and higher model revisions after errors (backward coupling changes) relative to the 60 ms tone consistent with models undergoing more revision in the second block than first block context.

Next, the difference in change in connectivity modulation associated with the deviant in the second block context (30 ms tone) was compared to that of the deviant in the first block context (60 ms tone) as a function of superordinate surprise or before versus after the change in block length. The right panel of Figure 3 therefore reflects the connections that were differentially affected by the interaction between local and superordinate surprise for a deviant in the second block context relative to first block context. As expected, violation of superordinate patterning was associated with significant differences in the changes in connectivity for the two tones which were marked by increased ascending connectivity at lower levels (A1-STG), decreased ascending connectivity at higher levels (STG-IFG) and greater descending connectivity generally for the second deviant when compared to the first deviant after the change in block length. Relative to the first-deviant (60 ms) tone, the second-deviant (30 ms) tone was additionally associated with reduced self-inhibition (increased gain) at bilateral A1 and decreased gain at bilateral STG after the superordinate change. These differences in connectivity change are consistent with greater influence of low-level prediction errors (A1 to STG) for the second deviant compared to the first deviant after superordinate patterning is violated, but a comparatively lower level of impact of higher-level prediction errors (STG to IFG) for the second deviant relative to the first deviant following this change. The observed connectivity changes could be considered consistent with the patterns of MMN amplitude modulation observed here (see Figure S1) and in previous AEP studies (34, 35), where the conflict between greater gain at lower levels (PE for local deviations) and greater PE suppression (descending predictions) throughout the hierarchy for the second-deviant tone might explain the smaller net MMN modulation for this tone than the first-deviant tone after the block length change. It is also consistent with the notion that after superordinate change, the occurrences of the first deviant have a more prominent influence over the remodelling of longer-term predictions reflected in the comparatively higher connectivity in forward connections between STG and IFG for the 60 ms tones.

## Discussion

The present study involved the novel application of DCM to examine brain responses elicited by the violation of hierarchical regularities in sound. This analysis revealed differential changes in connectivity in underlying brain networks before and after violations of local, intermediate and superordinate patterns during the sound sequence, and differed for the two tones based on their relative probability at sequence onset. These results provide further evidence that the brain is capable of unsupervised learning over multiple timescales simultaneously, and that prediction models are not a veridical representation of the local context but are modulated by higher-order representations. These findings are in concert with order effects previously observed in AEPs from scalp recordings, and give validity to the use of AEPs to study hierarchical inference and learning. Furthermore, the DCM supports a structural and functional architecture that is consistent with hierarchical predictive coding (18), and its neurobiological implementation within the canonical microcircuit model (55). In previous studies of the multiple-timescale paradigm, differential patterns of MMN modulation to the two tones were assumed to reflect how the updating of predictions in response to surprise at a given level will be constrained by internal models held at multiple levels of hierarchical inference. Here we will consider how each of these levels of surprise are substantiated within the hierarchical levels of the DCM, and more specifically how changes in predictions, prediction error and precision are reflected in changes in ascending, descending, and intrinsic connectivity respectively.

### Local deviance – Deviant relative to standard

Local surprise occurred at any given point throughout the sequence and was represented by the occurrence of the relatively less probable deviant among a series of relatively more probable standard tones, as in traditional oddball paradigms. Similar patterns of connectivity change were observed for deviants in both the first and second block context (i.e., both the 60 ms and 30 ms tone as deviant). The increase in inhibitory gain at rA1 and network-wide increase in ascending connectivity are consistent with previous modelling of deviant responses (e.g., (63, 64)) and are interpreted to reflect an increase in bottom-up prediction error signalling when a tone is deviant relative to standard. However, the increase in backward connectivity is surprising, and may reflect a true increase in the modulatory influence and/or remodelling of top-down predictions for a deviant relative to standard tone, or an absent decrease relative to standards. In either case it is possible that this anomalous result could reflect the impact of hierarchical learning, unique to the present study’s novel modelling of the multiple-timescale paradigm, on how prediction errors are modulated over the course of the overall sound sequence.

The deviant in the second block context (30 ms tone) was further associated with reduced self-inhibition (i.e., increased gain) at bilateral STG, consistent with increased prediction error signalling at higher levels required to increase new learning and override the suppression of PE associated with this tone having previously been redundant, with this reflected in the AEPs as MMN amplitude gradually increasing over time for the second deviant (34). These changes are consistent with a theoretical predictive coding treatment where deviance is assumed to trigger increased ascending error signalling and changes in the self-suppression of prediction errors in order to drive new learning and revise descending predictions, and are in keeping with previous DCM analyses of deviant versus standard tones in auditory oddball and roving standard paradigms (43, 44).

### Intermediate deviance – First vs Second deviant

The impact of higher-order surprise was investigated at two levels: intermediate surprise represented by changes in the specific tendencies of each tone between the original to reversed block type, and superordinate surprise represented by the change in block length defined by regular alternation in tone tendencies. Having established the connectivity changes associated with a deviant versus standard tone in general, we looked for distinct patterns of connectivity change in deviant responses before and after these specific points in the sequence, which might reflect how responses to local deviance are weighted by changes in the precision of higher order predictions when violated. The impact of intermediate surprise was investigated by comparing connectivity underlying the response to second deviants (30 ms tone), relative to first deviants (60 ms tone). Relative to the first deviant, second deviant responses were associated with increased descending coupling from bilateral IFG to STG, increased self-inhibition (decreased gain) at higher levels (STG and lIFG) and gain changes at A1 resembling that seen for a local deviant (decreased self-inhibition at lA1 and increased at rA1).

These patterns of difference in the connectivity changes are largely consistent with the theory that the precision of prediction errors, and subsequently the rate of new learning, were lower in the second block context where the 30 ms tone is deviant. Lower precision is evident in the increased strength of descending connections and decreased gain on superficial pyramidal cells at higher levels reflecting the greater influence of descending predictions (i.e. deviant occurrence leading to model updating) and stronger suppression of ascending prediction errors for this initially redundant tone compared to the 60 ms tone which was initially deviant. Meanwhile, the gain changes at bilateral A1 and increased ascending connections resembling that seen for a local deviant are consistent with the idea that this tone is still recognised as a new deviant, however is likely associated with a slower learning rate within this block due to lower precision. This interpretation is consistent with previous AEP studies of the multiple-timescale sequence demonstrating a gradual increase in MMN amplitude within blocks to the second deviant only, presumably due to a reduced rate of learning about this previously redundant tone as deviant (e.g., 38).

### Superordinate deviance – Before versus after a block length change

The final, superordinate level of surprise inherent in the sound sequence involved violation of the regular rate at which tone probabilities change (i.e., every 0.8 min in unstable components and every 2.4 min in stable components). Learning about this regularity occurred over the longest timescale and was violated only twice within the paradigm– once in the transition from unstable to stable components (increasing-stability sequence), and once in the transition from stable to unstable components (decreasing-stability sequence). When comparing the connectivity associated with each tone as deviant after this superordinate violation, the second context deviant (30 ms tone) was associated with lower ascending connectivity to IFG, greater descending connections, increased gain at A1 and decreased gain at STG relative to the first context deviant (60 ms tone). These differences are consistent with first deviant errors being treated as more informative and the first-deviant context being assigned high model precision, leading to a more significant impact on precision weightings for this tone when higher order patterning is violated. When the block length is violated, the assumptions underlying the block length predictions must be revisited (the model must be updated) and these are purported to be represented in higher levels of the network (18). The comparatively lower level of model precision (less descending influence) for the first context deviant after higher order patterning is broken, alongside the increase in gain and forward error signalling to higher levels (STG-IFG) and lower gain of PE associated with local deviations for the first context deviant tone after the superordinate change are all consistent with remodelling of higher order predictions based on this tone. This result supports the hypothesised role of higher hierarchical levels in longer term pattern learning and an adjustment to the weighting of this influence when these more global patterns are violated.

The Bayesian brain hypothesis purports that learning rates are dynamically adjusted to best minimise surprise by differentially weighting prediction errors at various levels to distinguish reliable changes from random fluctuations in the world. We propose that this accounts for the differential patterns of AEP amplitude and associated connectivity changes observed to the two tones throughout the sequence. Namely, that in the absence of any existing priors, high precision is afforded to the binary categorisation of the two tones as more or less probable at sequence onset (local surprise), leading to a lower accumulation of precision in the new tendency of these tones (intermediate surprise) after tone probabilities change. This is observed as a lower precision of prediction errors associated with the previously redundant tone in the second context through decreased gain on deviant response relative to the first context. Similarly, a more distributed pattern of disinhibition of superficial pyramidal cells is seen for a second context deviant following local surprise compared to a first context deviant, likely reflecting the increased learning rate required to override prior learning of the 30 ms tone as uninformative in the first context. Meanwhile, superordinate surprise requires the revision of beliefs about the general volatility of tone probabilities, which similarly shows lower precision associated with the second context deviant through less marked gain modulation, and higher influence of descending connectivity compared to the initial deviant (60 ms) tone specifically after the superordinate change.

In previous studies, differential modulations of MMN amplitude to the first and second deviant tone have persisted across as many as four repetitions of the same sound sequences (36), and are shown only to be altered when superordinate patterning is abolished or violated (35, 65) or if the participant has prior experience with the sounds (48) or prior knowledge of the sequence structure (66). The apparent failure to override a first-impression, even after four repetitions of the same sound sequence, implies that the system maintains differential precision weightings for the two deviants unless a substantive level of surprise is encountered (36). In the present data, these assumptions are reflected in a higher backward coupling strength for the second-deviant 30 ms tone compared to the first-deviant 60 ms tone overall, generally lower precision of the 30 ms than 60 ms deviant, and the precision of prediction errors becoming more precise for 30 ms than 60 ms deviants at lower levels after the superordinate change (at A1), but comparatively less precise than 60 ms deviants at the higher levels (STG) after the higher-order pattern violation when block lengths change.

The results also lend further support to the assertion that perceptual inference engages a hierarchical network architecture where more rostral projections such as those to the prefrontal cortex should be responsible for generating and updating beliefs about longer-term patterns whilst more caudal regions are sensitive to short-timescale change (18). The DCM demonstrated that selective deviant-specific disruptions in these more rostral projections from STG to PFC were primarily seen when the superordinate pattern was violated. These data are consistent with the notion that superordinate patterns must be present and learned for the effect to be seen given previous observation that order-effects are abolished when no superordinate patterning is available (65) or the participant (and therefore the prefrontal cortex and selective attention) is otherwise engaged by a cognitively demanding visual task (66).

In the present study we have drawn exclusively on AEPs as a vehicle through which to study perceptual inference and learning, with extension to underlying network connectivity through the application of DCM. The auditory MMN and AEPs more generally are an ideal candidate for tracking this type of statistical learning given the importance of temporal regularity in audition (e.g., to decipher grammar and semantics in human language), the ease at which the statistics of sound sequences can be manipulated, and the established sensitivity of the AEP to changes in these statistics. AEPs further confer an ease of measurement through their accessibility, automaticity and simplicity of the paradigms through which they are elicited, and are suited to the study of more difficult populations such as clinical groups, infants and the elderly. Traditional analyses of the AEP are often restricted to chosen electrodes and latencies based on the component of interest and draw interpretations about neural activity as observed at the scalp. DCM, in contrast, is applied to the entire time-course and sensor space of the epoch and is more sensitive in generating a biologically informed mathematical models of neural activity below the scalp. Whilst DCM has previously proven useful in elucidating the auditory MMN in an oddball sequence, this study is the first to our knowledge to implement DCM of AEPs within a nested hierarchical sequence. Our results confirmed hypotheses about underlying neural mechanisms derived from scalp-level AEP data (32,34,35,38,48,65,66), and demonstrate the utility of DCM to extend on the use of AEPs as a proxy for learning processes within multiple timescale paradigms.

The two methods are different in that DCM analysis captures the entire epoch where previous AEP analyses of the multiple timescale paradigms have restricted analysis to the 90-210 ms window surrounding the MMN. DCM is not limited to a priori components, but rather is sensitive to the contribution of any number of different AEP components occurring up to 300 ms post-stimulus (e.g., P3a) to the observed connectivity change. The changes presented, despite reflecting activity across the entire epoch, remain consistent with interpretations offered for the patterns shown previously in AEPs extracted from a smaller sampling window representative of the MMN, suggesting that the impact of additional components to the observed order effects are minimal, or are subject to the same patterns of order-driven modulation as seen for the MMN.

A limitation of the DCM is in model selection, where relative evidence is calculated only for the model space which is pre-defined. This entails the possibility that the data could be more accurately explained by an alternative model which is not captured in the model space. The chosen model space represents that which is to our knowledge best supported by the literature pertaining to auditory change detection and MMN generation (43–45). We are confident it represents the best estimate of a plausible model for generation of the observed responses based on current knowledge and also, how connectivity changes within a commonly accepted inferential network structure can account for our data. Additional assumptions were made in the process of source selection and again represent best-practice estimation through the use of MNI coordinates consistent with previous DCM studies (43,44,46). A further complication of DCM is that interpreting the complex and nonlinear dynamics of the brain it is designed to capture is not straightforward, as a number of interactions between underlying subpopulations could give rise to the observed connectivity change (e.g., see (43, 67) for discussion). There could therefore be many causes of the observed connectivity changes, however we have suggested the most likely interpretation that could be drawn based on previous studies in concert with the present data.

Whilst the present study is also limited in its focus on applying DCM to AEPs, recent trial-by-trial analyses have supported Bayesian learning across multiple levels of volatility during a multi-feature visual roving standard paradigm (68), and similar models of hierarchical Bayesian belief updating have also been successfully applied in DCM studies of visuospatial attention (69). The current findings could therefore be interpreted as supporting hierarchical Bayesian inference as a general framework for multi-modal inference and learning in the brain, rather than merely a specific feature of auditory processing. The current results lend support to hierarchical models of Bayesian learning, however may also have parallels to the Hierarchical Gaussian Filter (HGF), a recently formulated generative model of perceptual learning which employs Bayesian principles to trial-by-trial data (20,39,68). The HGF estimates hidden states from limited sensory input using hierarchical belief updating over multiple levels which are representative of the varying degrees of volatility in the environment. For example, the first level represents overall beliefs about possible states, the second level represents the current belief in the relative probability of encountering each state, and a third level represents the likelihood that these probabilities will change – levels which could be considered analogous to the various degrees of patterning in the present study’s sound sequence (see (65) for further discussion). Future studies may therefore look to directly embed the HGF model within DCM in order to provide conclusive empirical support for this form of learning within neural circuitry. Further, applications of DCM can contribute to more specific predictions about functional architecture and neurobiology given their increased sensitivity to mechanisms underlying the responses observed at the sensor level (42). Elucidating the neurophysiological basis for perceptual inference and learning in the healthy brain has further been considered to have important implications for understanding the etiology of disorders such as schizophrenia where these inference processes differ significantly (e.g., 66,67).

## Materials and Methods

### Participants

Participants were 19 healthy adults (15 female; aged 18-53 years, *M* = 25.26 years, *SD* = 11.44 years) recruited from undergraduate psychology students at the University of Newcastle and community volunteers. Exclusion criteria included current diagnosis of, or treatment for, a mental disorder per *Diagnostic and Statistical Manual of Mental Disorders – Fifth Edition* (47) criteria, history of head injury or neurological disorder, hearing loss, regular recreational drug use, heavy alcohol use or a first-degree relative with schizophrenia.

### Ethics Statement

All participants provided written informed consent to participate in the study protocol as approved by the University of Newcastle Human Research Ethics Committee prior to participating (approval number H-2012-0270). Reimbursement was provided in the form of course credit for students or gift vouchers for volunteers as compensation for time and expenses incurred.

### Stimuli and Sequences

Sound sequences were arranged as outlined by Fitzgerald and Todd (34), and consisted of 1000 Hz pure tones, presented over binaural Sennheiser HD280pro headphones at 75 dB with a 300 ms stimulus onset asynchrony. Sounds were a 30 ms and 60 ms tone created with a 5ms rise/fall time and a 20 ms and 50 ms pedestal respectively. Sound sequences consisted of the two tones alternating in the role of repetitive standard (*p* = .875) and rare deviant (*p* = .125) across blocks at two different regular rates to form a stable and relatively unstable sequence component, which were further arranged to create one increasing-stability and one decreasing-stability sequence. The general arrangement of sound sequences was used in a number of previous studies of auditory processing across multiple timescales (32,35–38,48), with the specific variation used by Fitzgerald and Todd (34) depicted in Figure 4.

**Figure 4.**
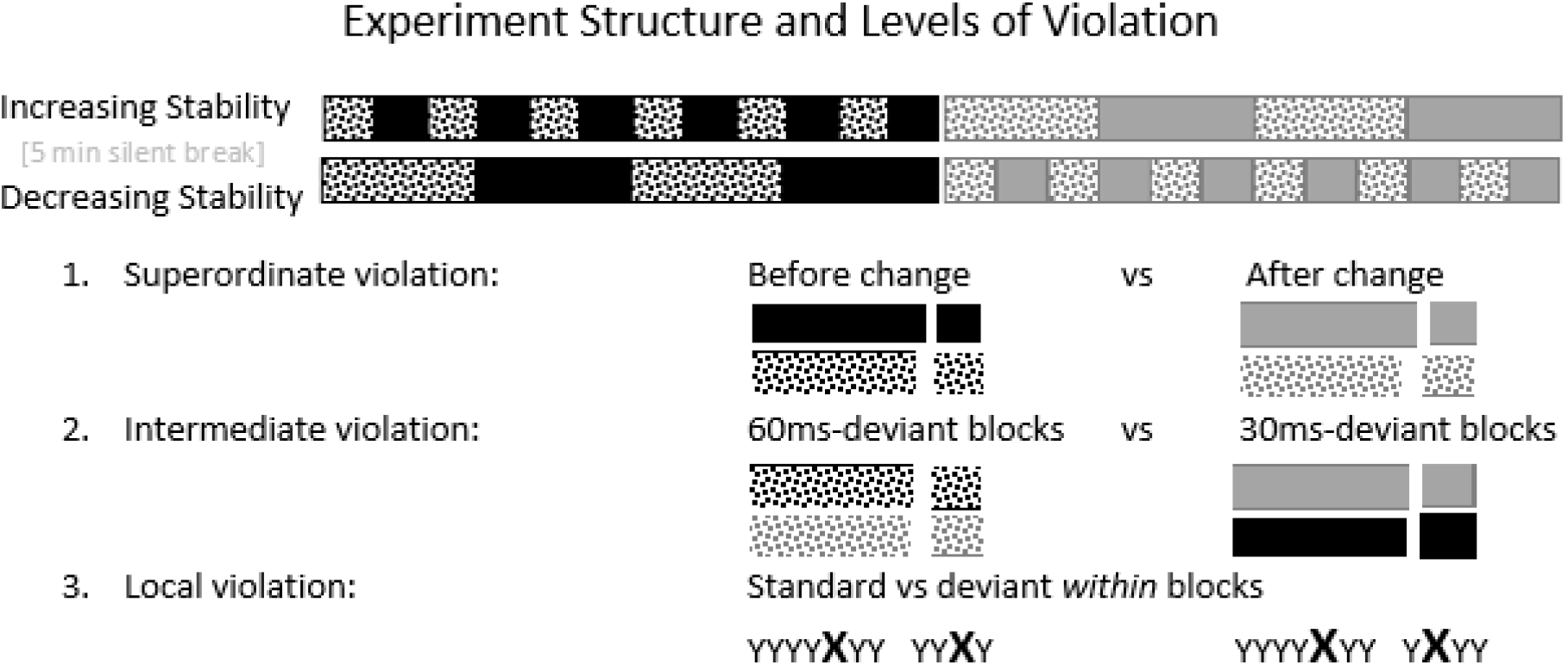
Representation of sound sequence structure and levels of pattern violation nested within the sound sequence. Local violation occurred within all blocks by presentation of a rare deviant (*p* = .125; represented as X) amongst a series of repetitive standards (*p* = .875; represented as Y), intermediate violation occurred following the change in contexts where tone probabilities alternated from a 60 ms deviant and 30 ms standard (first-context block) to a 30 ms deviant and 60 ms standard (second-context block; spotted vs solid blocks), and superordinate violation occurred following a change in block length in each of the sound sequences (black vs grey blocks).

Within these sound sequences, local, intermediate and superordinate regularity violations were represented by low-probability deviants, change in tone probabilities between the first and second block contexts (solid vs spotted blocks in Figure 1), and change in block lengths (black vs grey blocks in Figure 1) respectively. Importantly, each stable component consisted of 4 blocks of 480 tones and each unstable component consisted of 12 blocks of 180 tones – each adding to a total of 9.6 mins duration, 1960 tones in total and an equivalent number of standard and deviant tones overall. The only difference between sequence components was the maximum period of time over which tone roles remained stable. All participants heard the two sequences in the same order as depicted in Figure 1 – an “increasing-stability” arrangement consisting of the unstable followed by stable component, before a “decreasing-stability” arrangement consisting of the stable followed by unstable component. The two sequences were separated by a 5 minute silent break where participants were permitted to move to minimise discomfort and for the predictive system to “reset”. Previous multiple-timescale sequences have demonstrated no differences in order-driven effects between counterbalanced sequences when separated by a 2-5 minute break (31).

### EEG data collection and pre-processing

Fitzgerald and Todd (34) obtained a continuous electroencephalography (EEG) recording during presentation of the sound sequences via a SynAmps2 Neuroscience© system using a 1000 Hz sampling rate, high-pass 0.1 Hz, low-pass 70 Hz, notch filter 50 Hz and fixed gain of 2010. The EEG setup consisted of 64 electrodes in accordance with the International 10 ± 10 system with Modified Combinatorial Nomenclature (49) and included one electrode at the nose and each of the bilateral mastoids for use as reference. Additional electro-oculogram electrodes were placed 1cm from the outer canthi of each eye, and directly above and below the left eye to monitor eye movements. Impedances were reduced to below 5kΩ prior to recording.

The continuous EEG recordings from Fitzgerald and Todd (34) were re-processed using Neuroscan Edit^©^ software for suitability to the current DCM analysis. Adjustments involved band-pass filtering to a range of 0.5 to 40 Hz with 12dB drop-off and zero phase. Manual artefact rejection and bad channel exclusions were carried over from the previous analysis. Eye blink corrections were also completed in the previous analysis using a EEG-VEOG covariance analysis, linear regression and point-by-point subtraction procedure (50). Data was epoched from 50 ms pre-stimulus to 300 ms post-stimulus, and any epochs containing frequencies exceeding ±70 μV discarded prior to averaging.

All subsequent processing steps were undertaken using Statistical Parametric Mapping (SPM) software (version 12, revision 6906). SPM is a freely available academic software package specialised for the spatially extended statistical analysis of brain imaging data and which is suitable for DCM analyses (51).

Data were common average referenced as recommended for the application of DCM of EEG data (52) and a revised baseline correction from 25 ms pre-stimulus to 25 ms post-stimulus was applied to improve the existing baseline for present purposes. Single-subject and grand averages were subsequently generated for the response to each tone (60 ms, 30 ms) as standard and deviant in each sequence component (stable, unstable) and structure (increasing-stability, decreasing-stability) resulting in 16 grand averages in total.

Data was re-organised for analysis to specifically test the effects of local, intermediate and superordinate pattern violations by comparing the same tone as standard versus deviant (local violation), the first-context versus second-context deviant (i.e., 60 ms vs 30 ms tone; intermediate violation), and the two tones as deviants before versus after a change in block length (superordinate violation) respectively. Block length (stable, unstable) was not a factor in the current analysis, given that a change in block length from stable to unstable sequence components produced the same MMN modulation patterns to a change from unstable to stable (i.e., the changes were tied to a change in block length rather than specifically related to block length, (34). Since we focused only on robust effects in the data for subsequent DCM analysis, order was therefore collapsed over block length, with first-heard components (unstable in increasing and stable in decreasing) compared to second-heard (stable in increasing and unstable in decreasing) components (i.e., the sequence components differentiated by black vs grey colouring in Figure 1). The remaining sections will refer exclusively to factors of role (standard, deviant), tone (30 ms, 60 ms) and order (heard-first, heard-second) accordingly. Analysis by role represents local violations by comparing standard vs deviant, analysis by tone represents intermediate violations by comparing the two deviant types, and analysis by order represents superordinate violations by comparing the two global sequence structures.

### Sensor space analyses

The common-average referenced ERP waveforms were analysed to confirm the presence of the same characteristic patterns previously observed in the mastoid-referenced data by Fitzgerald and Todd (34). To assess for sensitivity to deviant tones and changes in the tendency of these tones we compared ERPs to standards and deviants separately for each tone (30 ms vs. 60 ms), using family-wise-error corrected paired *t*-tests to test for significant (*p* < .05) differences in these responses at each sampling point within the epoch (corrected over sampling points). For statistical analyses of ERP time-series, data were extracted from the F4 channel given that both MMN and previous observations of order-driven effects have been shown to be frontal and right-hemisphere maximal (e.g., 34,53,54). Sensitivity to higher-order pattern violations was investigated by family-wise-error corrected *t*-tests investigating for significant (*p* < .05) differences in the ERP to each tone as deviant before and after a change in block length in each condition over the entire epoch at F4.

### Dynamic causal modelling

Further analysis was undertaken using DCM to estimate population output and connectivity parameters associated with the three key patterns observed in AEP data: sensitivity to tone probabilities (local violation), changes in the probabilities of the two tones (intermediate violation), and changes in the volatility of these probabilities (superordinate block-length violation). DCM allows for a mapping from data measured at the sensor level to source-level activity, in a sparse network of interconnected sources, each consisting of a set of neural populations based on a canonical microcircuit architecture (55). The activity in each source evolves as described using coupled differential equations which model the dynamics of postsynaptic voltage and current in each neural population. These populations (spiny stellate cells, superficial and deep pyramidal cells, and inhibitory interneurons) have distinct connectivity profiles of ascending and descending projections linking different sources (extrinsic connectivity) and coupling neural populations within each source (intrinsic connectivity). DCM based on canonical microcircuits has been used in several other studies of mismatch responses (e.g., 56,57), and validated using invasive recordings in humans (58).

Model inversion in DCM is susceptible to local maxima issues due to the inherently non-linear nature. To overcome this potential issue we implemented Parametric Empirical Bayes (PEB), an iterative hierarchical implementation of the empirical Bayesian inversion method (59) where group-level effects are inferred by fitting the same model to each participant’s data under group constraints (e.g., the assumption that model parameters are normally distributed in the participant sample) updating the posterior distribution of the individual DCMs and re-inverting the model over several iterations. This process was applied using the built-in SPM 12 function spm_dcm_peb_fit.m.

The DCM adopted a standard electromagnetic forward model based on the Boundary Elements Model (BEM) in Montreal Neurological Institute space as the default SPM 12 template (52). Lead-fields specified by the forward model were used to reconstruct AEP responses at all electrodes and latencies (0-300 ms) from six cortical sources considered for inclusion in the DCM: bilateral primary auditory cortex (A1), bilateral superior temporal gyrus (STG) and bilateral inferior frontal gyrus (IFG), using the following MNI coordinates (43, 46): left A1 [-42, −22,7], right A1 [46, −14, 8], left STG [-61, −32, 8], right STG [59, −25, 8], left IFG [-46, 20, 8], right IFG [46, 20, 8]. Changes in extrinsic (between cortical sources) or intrinsic (within cortical sources) connections were quantified as model parameters giving rise to differences between AEPs. The free-energy approximation to model evidence was used as a metric of model fit to the data, penalised by model complexity.

DCM analyses were conducted in two steps. The first analysis modelled changes in standard and deviant responses to assess for the expected effects of deviance on connectivity parameters (e.g., ascending connections) and confirm the validity of this application of DCM to data extracted from the multiple-timescale paradigm, whilst also modelling the effect of local violation represented by a deviant relative to standard tone. The second analysis modelled changes in the deviant AEP only given that order-driven effects on MMN amplitude modulations are driven primarily by the deviant AEP (see Results), and focused on differential changes in connectivity associated with intermediate violations (second/30 ms vs first/60 ms deviant) and superordinate violations (before vs after a change in block length). Given that the modulations of interest are based on order rather than tone properties, in the first analysis responses to the second deviant (30 ms) and first deviant (60 ms) tone were modelled separately to give a pure measure of network changes to a deviant which were not conflated with the task of explaining variance due to order effects.

Individual model inversion was conducted fitting separate models to the two levels of intermediate deviance (30 ms as deviant vs. 60 ms as deviant) over two factors - local deviance (standard versus deviant), superordinate deviance (before and after a change in block length) - and their interaction. In the second analysis tone type was included as a factor within a single DCM to permit the direct contrast of how order driven effects differentially impact connectivity underlying responses to the two tones after a superordinate change. Here individual model inversion was conducted fitting the single DCM over the two factors of intermediate deviance (second deviant/30 ms tone vs.first deviant/60 ms tone) and superordinate deviance (heard-first versus heard-second), and their interaction. In both analyses individual model inversion was applied using empirical priors over the six-source network permitting changes in ascending, descending and intrinsic gain parameters.

Bayesian model reduction (BMR) was used to identify the parameter changes that best explained the observed AEP data and estimate the variation in these parameters caused by local, intermediate and superordinate deviations in the respective analyses. BMR uses inversion of the “full” model incorporating changes in all identified parameters to estimate model evidence for a range of “reduced” models where some parameters are not permitted to vary (60). The chosen model space in both analyses examined each combination of changes in ascending connections, descending connections and modulatory gain parameters for each factor, resulting in 8×8 factorial models (ascending, descending, ascending/descending and null, each with and without modulatory gain changes).

The winning models were next entered into PEB to hierarchically estimate the variation in parameters which explained systematic changes in response to each factor, comparing a standard relative to deviant in the first analysis, second deviant (30 ms) relative to first deviant (60 ms) tone in the second analysis, and first-heard relative to second-heard sequence component in both analyses. This approach permits estimation of a general linear model for model parameters across individually inverted (first-level) DCMs. Regressors in the second level model included the group mean for each factor (local and superordinate deviance in the first analysis, intermediate and superordinate deviance in the second analysis) and random subject effects. In the first analysis intermediate deviance formed an additional second-level regressor given that the 30 ms and 60 ms tone were modelled separately for each participant. BMR was subsequently applied to identify significant changes in parameters due to these second level factors. Parameters at the second level were derived from the canonical microcircuit model for DCM and included baseline estimates describing extrinsic connections (between sources; A), intrinsic connections (between neural populations within cortical sources; G), and activity-dependent effects on intrinsic connections (modelled as activity-dependent superficial pyramidal cell self-inhibition; M), as well as modulatory parameters describing the effects of the experimental manipulations on extrinsic and intrinsic connections (B) and the activity-dependent effects on intrinsic connections (N). The BMR generated outputs of estimated free-energy approximation to the log-evidence for each second level model (used to compare models and select the winning model), the parameter changes associated with local and superordinate violations, and a Bayesian 95% confidence interval for each as a measure of uncertainty in the estimates. Significance testing followed a recently developed procedure for empirical Bayes that is considered more robust to repeated testing than t-tests and involves estimating the proportion of the probability distribution that falls either side of zero for each parameter against a statistical threshold of 0.995 (61).

## Acknowledgements

KF acknowledges receipt of an Australian Government Research Training Program (RTP) Scholarship. We thank Gavin Cooper for his assistance in programming these experiments and Professor Michael Breakspear for comments on an early version. This research was supported by funds provided by the National Health and Medical Research Council of Australia (APP1002995).

## Supporting Information Legends

Figure S1. SPM output highlighting significant spatio-temporal increases (F-test, p < .05 family-wise error-corrected) in deviant relative to standard responses across 2D sensor space and time (left panels) and at peak (right panel) for the 30 ms and 60 ms tone.

Figure S2. Grand average difference waveforms at F4 (where the MMN is maximal) for the second deviant/30 ms (black lines) and first deviant/60 ms tone (grey lines) when heard before (solid lines) and after (broken lines) a change in block length in unstable (left panel) and stable (right panel) sequence components. Shaded bars indicate the latency windows over which mean amplitudes were extracted for quantification of the MMN (160-180 ms) and P3 (240-260 ms) components, based on a 20ms window capturing the common peak for a majority of individual averages across conditions. Significant differences in mean amplitude are indicated by an asterisk (*p* < .05).

Figure S3. Grand average AEPs to the 30 ms (black lines) and 60 ms (grey lines) tone as deviant (A; top panel) and standard (B; bottom panel) when heard before (solid lines) and after (dashed lines) a change in block length for unstable and stable sequence components. Horizontal bars in (B) indicate periods of significant difference between standard and deviant waveforms elicited to the 30 ms (black) and 60 ms (grey) tones across orders, and periods of significant difference between heard-first and heard-second components in the deviant response for the 60 ms tone (blue; one-tailed *t*-test, *p* < .05, using the procedure outlined by (62) for statistical analysis of EEG waveforms). There were no periods of significant difference in the deviant waveforms for a 30 ms tone when heard first versus heard second.

**Table S1.** Estimated changes in forward, backward and intrinsic connections within/between sources associated with a 30 ms tone when encountered with deviant probability as compared to when encountered as a standard. Significant changes are marked by an asterisk, and were assessed as 99.95% of the estimated probability distribution demonstrating a change greater or less than zero.

|  | Forward | Backward | Intrinsic |
| --- | --- | --- | --- |
| lA1-lSTG | 0.282* |  |  |
| rA1-rSTG | 0.880* |  |  |
| lSTG-lIFG | 0.865* |  |  |
| rSTG-rIFG | 0.852* |  |  |
| lSTG-lA1 |  | 1.211* |  |
| rSTG-rA1 |  | 0.061 |  |
| lIFG-lSTG |  | -0.075 |  |
| rIFG-rSTG |  | 0.176 |  |
| lA1 |  |  | -0.371* |
| rA1 |  |  | 0.586* |
| lSTG |  |  | -0.402* |
| rSTG |  |  | -0.745* |
| lIFG |  |  | 0.081 |
| rIFG |  |  | 0.653* |

**Table S2.** Estimated changes in forward, backward and intrinsic connections within/between sources associated with a 60 ms tone when encountered with deviant probability as compared to when encountered as a standard. Significant changes are marked by an asterisk, and were assessed as 99.95% of the estimated probability distribution demonstrating a change greater or less than zero.

|  | Forward | Backward | Intrinsic |
| --- | --- | --- | --- |
| lA1-lSTG | 1.135* |  |  |
| rA1-rSTG | 2.215* |  |  |
| lSTG-lIFG | 0.770* |  |  |
| rSTG-rIFG | 0.671* |  |  |
| lSTG-lA1 |  | -0.192 |  |
| rSTG-rA1 |  | 0.314* |  |
| lIFG-lSTG |  | 0.246 |  |
| rIFG-rSTG |  | < -0.001 |  |
| rA1 |  |  | -1.342* |
| lA1 |  |  | 0.462* |
| rSTG |  |  | -0.254 |
| lSTG |  |  | 0.210 |
| rIFG |  |  | -0.064 |
| lIFG |  |  | -0.004 |

**Table S3.** Estimated changes in forward, backward and intrinsic connections within/between sources associated with a 30 ms tone as deviant compared to a 60 ms tone as deviant. Significant changes are marked by an asterisk, and were assessed as 99.95% of the estimated probability distribution demonstrating a change greater or less than zero.

|  | Forward | Backward | Intrinsic |
| --- | --- | --- | --- |
| lA1-lSTG | -0.057 |  |  |
| rA1-rSTG | 0.470* |  |  |
| lSTG-lIFG | 0.033 |  |  |
| rSTG-rIFG | 1.11* |  |  |
| lSTG-lA1 |  | 0.010 |  |
| rSTG-rA1 |  | -0.216* |  |
| lIFG-lSTG |  | 1.168* |  |
| rIFG-rSTG |  | 0.181* |  |
| rA1 |  |  | -0.686* |
| lA1 |  |  | 0.144* |
| rSTG |  |  | 0.202* |
| lSTG |  |  | 0.250* |
| rIFG |  |  | 0.228 |
| lIFG |  |  | 0.147 |

**Table S4.** Estimated changes in forward, backward and intrinsic connections within/between sources associated with a 30 ms tone as deviant compared to a 60 ms tone as deviant before versus after the change in block length. Significant changes are marked by an asterisk, and were assessed as 99.95% of the estimated probability distribution demonstrating a change greater or less than zero.

|  | Forward | Backward | Intrinsic |
| --- | --- | --- | --- |
| lA1-lSTG | -0.018 |  |  |
| rA1-rSTG | 0.486* |  |  |
| lSTG-lIFG | -0.368* |  |  |
| rSTG-rIFG | -0.225* |  |  |
| lSTG-lA1 |  | 0.437* |  |
| rSTG-rA1 |  | 0.244* |  |
| lIFG-lSTG |  | 0.316* |  |
| rIFG-rSTG |  | -0.039 |  |
| rA1 |  |  | -0.702* |
| lA1 |  |  | -0.228* |
| rSTG |  |  | 0.186* |
| lSTG |  |  | 0.337* |
| rIFG |  |  | -0.303* |
| lIFG |  |  | 0.147 |

